# methyl-ATAC-seq measures DNA methylation at accessible chromatin

**DOI:** 10.1101/445486

**Authors:** R Spektor, ND Tippens, CA Mimoso, PD Soloway

## Abstract

Chromatin features are characterized by genome-wide assays for nucleosome location, protein binding sites, 3-dimensional interactions, and modifications to histones and DNA. For example, Assay for Transposase Accessible Chromatin sequencing (ATAC-seq) identifies nucleosome-depleted (open) chromatin, which harbors potentially active gene regulatory sequences; and bisulfite sequencing (BS-seq) quantifies DNA methylation. When two distinct chromatin features like these are assayed separately in populations of cells, it is impossible to determine, with certainty, where the features are coincident in the genome by simply overlaying datasets. Here we describe methyl-ATAC-seq (mATAC-seq), which implements modifications to ATAC-seq, including subjecting the output to BS-seq. Merging these assays into a single protocol identifies the locations of open chromatin, and reveals, unambiguously, the DNA methylation state of the underlying DNA. Such combinatorial methods eliminate the need to perform assays independently and infer where features are coincident.

## INTRODUCTION

Active promoters, enhancers, and other gene regulatory sequences are typically bound by sequence-specific transcription factors (TFs), free of nucleosomes, and these facilitate transcription. Such regulatory sequences can be identified by methods that detect nucleosome-depleted regions (NDRs), including DNase-seq, which identifies NDRs by their hypersensitivity to DNase I (Thurman et al. 2012); FAIRE-seq, which identifies NDRs according to their reduced protein content (Gaulton et al. 2010); and ATAC-seq, which identifies NDRs based on their increased accessibility to Tn5 transposase integration, and accordingly are called Transposase hypersensitive sites (THS) (Buenrostro et al. 2013). There is considerable agreement among the regions identified by each assay. ATAC-seq has received further use recently owing to its simplified workflow, reduced material requirements and lower background signals. Additional advancements such as Omni-ATAC (Corces et al. 2017) and Fast-ATAC (Corces et al. 2016) have further improved the utility of ATAC-seq.

DNA within NDRs may have different modification states, including methylation at the fifth carbon of Cytosine (5mC), and oxidized derivatives. In the mammalian genome, most 5mC is found at CpG dinucleotides, and is generally associated with transcriptionally inactive regions. Bisulfite sequencing (BS-seq) uses selective chemical deamination of unmodified cytosines to uracil, leaving 5mC unchanged. The extent of methylation at a given CpG in a sample is detected after amplification, sequencing, aligning reads to the genome, and then assessing the proportion of aligned reads that retained a C at a CpG, diagnostic of methylation, *vs.* a T, which reports an unmethylated residue.

Two features of BS-seq dramatically increase costs compared to routine sequencing assays. First, bisulfite treatment reduces the yield and complexity of DNA libraries, resulting in fewer reads uniquely aligning to the genome. Second, to reliably quantify the extent of methylation of a given CpG requires high read coverage. For these reasons, Reduced Representation Bisulfite Sequencing (RRBS) (Meissner et al. 2005) and derivatives (Boyle et al. 2012; Chatterjee et al. 2012; Garrett-Bakelman et al. 2015) have been used to focus analysis on CpG dense regions. However, not all gene regulatory sequences are detected by RRBS, and many regions that are detected are not regulatory.

Integrating results from assays for distinct chromatin features have defined novel categories of regulatory elements. These include bivalent promoters (Bernstein et al. 2006), enhancers (Heintzman et al. 2009), and widely observed chromatin states likely to harbor shared regulatory functions (Roadmap Epigenomics et al. 2015). In most of these studies, results from assays for single features are superimposed, and when a given locus has signals for multiple features, the features are inferred to be coincident on the same molecule. Though many inferences might be accurate, there is uncertainty inherent in such approaches, owing to the fact that samples commonly contain multiple sub-populations of cells, each with a characteristic chromatin state. Accordingly, the population-averaged results might report chromatin states found in no individual subpopulation of cells. Methods that combine assays for multiple chromatin features in a single protocol can eliminate this ambiguity for the features assayed. Here, we describe methyl-ATAC-seq (mATAC-Seq), a modification of ATAC-seq that combines ATAC-seq with BS-seq, identifying the locations of open chromatin, and the methylation state of the underlying DNA. In addition to providing more reliable assignments of chromatin states, mATAC-seq can focus DNA methylation analyses to accessible regulatory regions of the genome.

## RESULTS

Fig. 1 shows the workflow and sample results for mATAC-seq. It includes two primary modifications during the transposition step of the Omni-ATAC-seq protocol: (1), methylated oligonucleotides are loaded onto Tn5 to generate the transposome (Fig. 1A) which is then used to perform ATAC-seq (Fig. 1B); and (2), 5-methyldeoxycytosine triphosphate (5-mdCTP) is substituted for dCTP during the subsequent end repair step (Fig. 1C). These modifications protect the Nextera adapter sequences during the final step of mATAC-seq library preparation, bisulfite treatment of the tagmented DNA (Fig. 1D). Use of methylated oligonucleotides, and 5-mdCTP during end repair protects cytosines in the adaptors from deamination caused by bisulfite treatment, which is necessary for successful PCR amplification and sequencing of the resulting libraries. Sequenced libraries provide information on both DNA methylation and Transposase hypersensitivity (Fig. 1E).

**Figure 1:**
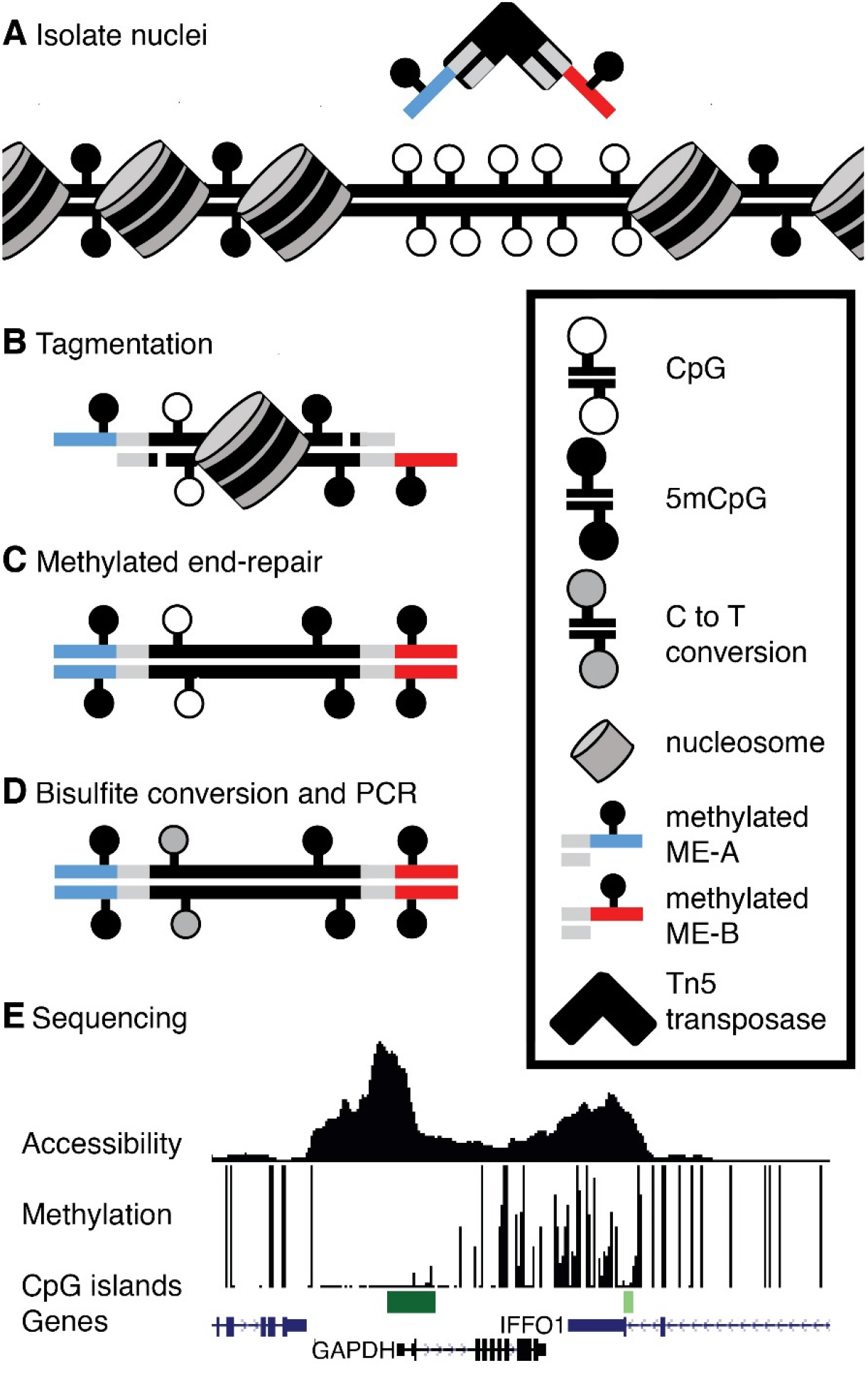
Overview of mATAC-seq; (A) Tn5 carrying methylated oligonucleotides (red and blue segments) is used to (B) perform tagmentation on nuclei at THS sites. (C) Tagmented DNA is end-repaired using 5mdCTP + dDTPs, purified, (D) Bisulfite converted, amplified, and (E) sequenced to measure DNA methylation and accessibility simultaneously; sample data are shown for one region in HCT116 cells. Peak height in accessibility track is proportional to read abundance; bar height in methylation track is proportional to extent of methylation at CpGs.

We applied mATAC-seq to nuclei prepared from HCT116 colorectal carcinoma cells. mATAC-seq reads in peaks were highly reproducible in biological replicates (r^2^=0.90, Fig. S1A). To validate that mATAC-seq captured open chromatin domains as well as conventional methods, we compared Transposase Hypersensitive (THS) sites found by mATAC-seq with those we identified using the standard Omni-ATAC-seq protocol (Fig. 2A-D) (Corces et al. 2017). Approximately 92% of called peaks found by Omni-ATAC-seq were found by mATAC-seq (Fig. 2A). There was also strong concordance between mATAC-seq and Omni-ATAC-seq with respect to gene features detected by both assays, with promoter regions being the most commonly identified features (Fig. 2B). In addition, reads in peaks identified by Omni-ATAC-seq and mATAC-seq were well correlated (Fig. S1A, B). Regions of greatest divergence include difficult to map regions such as repetitive elements, low complexity sequences, and simple repeat annotations (Fig. S1C). These analyses demonstrate that mATAC-seq detects open chromatin comparably to traditional Omni-ATAC-seq, and that protocol modifications that enable subsequent bisulfite sequencing do not compromise detection of open chromatin.

**Figure 2:**
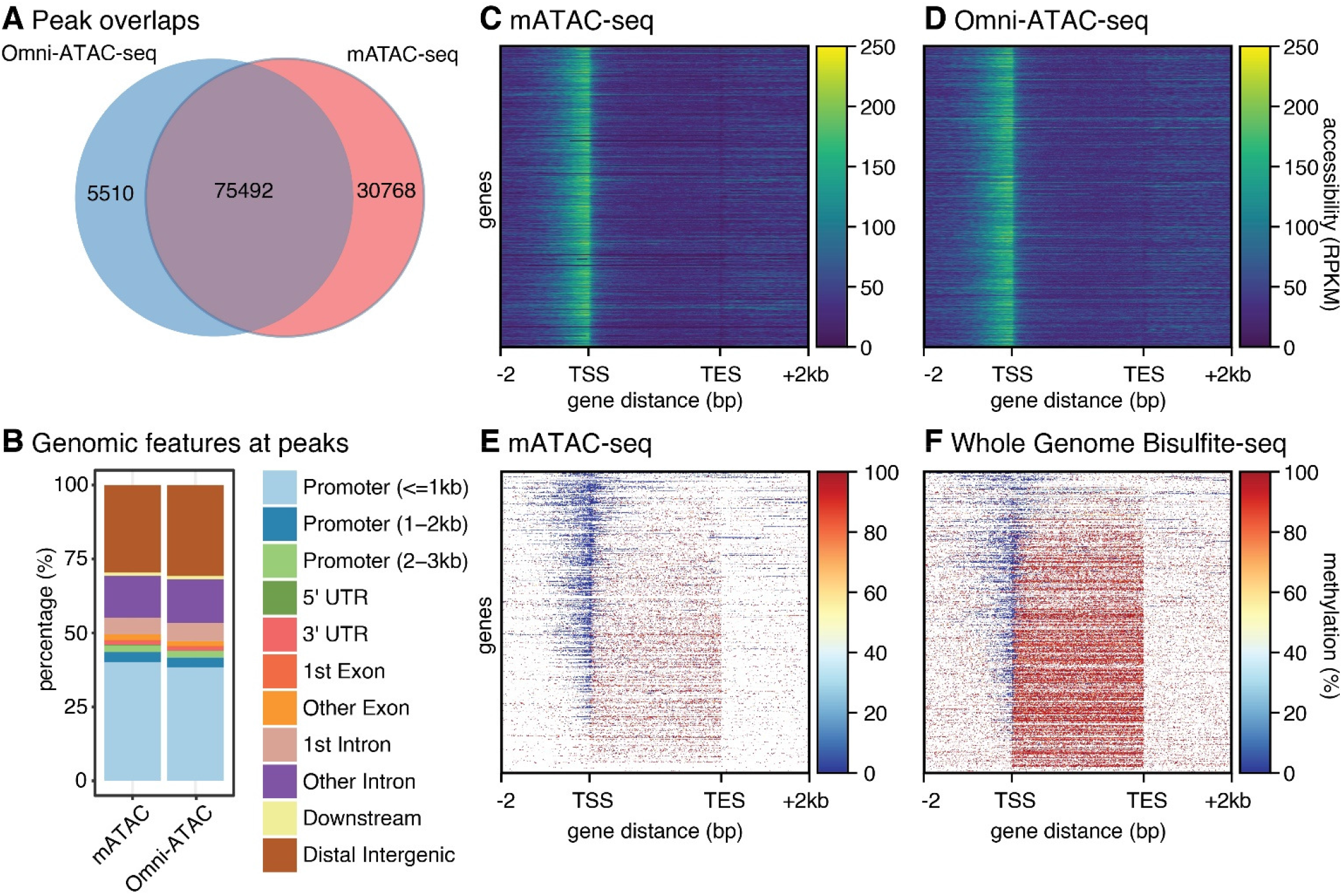
Comparison of methods; (A) Omni-ATAC and mATAC share a majority of peaks (B) Features at peaks are similar for mATAC and Omni-ATAC. (C) Accessibility in mATAC-seq is comparable to (D) Omni-ATAC-seq at gene bodies +/− 2kb, n=21,305. (E) methylation reported by mATAC is comparable to (F) WGBS at gene bodies +/− 2kb, n=21,305, though WGBS includes data absent from mATAC.

To validate that mATAC-seq identified DNA methylation patterns as reliably as conventional methods, we next compared the mATAC-seq methylation data with whole genome bisulfite sequencing (WGBS) data reported for HCT116 cells at THS sites and CpG islands (Blattler et al. 2014). DNA methylation detected by mATAC-seq replicates was highly reproducible at peaks (r^2^=0.83) and CpG Islands (r^2^=0.95) (Fig. S2A, B); and methylation levels reported by mATAC-seq correlated well with levels reported by WGBS at peaks (r^2^=0.68) and CpG Islands (r^2^=0.85) (Fig S2A, B). THS peaks identified by mATAC-Seq in HCT116 were predominantly unmethylated, and this agrees with existing WGBS data (Fig S2C, D). Fig. 2E and Fig. 2F report DNA methylation patterns assayed respectively by mATAC-seq and WGBS across gene bodies spanning from 2kb 5’ of transcriptional start sites (TSS) to 2kb 3’ of transcriptional end sites (TES). These patterns are consistent with the high correlations described above. We find these high correlations despite the fact that the assays were performed by different labs; also, WGBS and mATAC-seq assays are different in that mATAC-seq queries DNA methylation at open chromatin, whereas WGBS assays the entire genome, regardless of chromatin state. Our mATAC-seq data showed a reciprocal relationship between accessibility and 5mC density. These are in agreement with previous results from NOMe-seq (Kelly et al. 2012), which can also report sites of accessible chromatin and DNA methylation states but requires much greater sequencing depth. Both assays reveal that highly accessible chromatin is depleted of 5mC, and that there is an abundance of methylation in less accessible chromatin over gene bodies (Fig. 2E, F). Having shown that sites of open chromatin and DNA methylation states reported by mATAC-seq, Omni-ATAC-seq, and WGBS are in agreement, we concluded that mATAC-seq can be used to simultaneously identify the locations of the genome with accessible chromatin, and the methylation state of the underlying DNA. Because mATAC-seq measures accessibility and methylation in a single assay, it eliminates the inherent uncertainty about coincidence of chromatin features that can arise when ATAC and bisulfite assays are performed independently, and inferences are made after overlaying the two datasets, and at lower costs.

We extended our analyses of HCT116 cells, performing mATAC-seq on HCT116-derived *DNMT1* and *DNMT3B* double knock-out cells (DKO) (Rhee et al. 2002) to assess the functional significance of these methyltransferases on chromatin accessibility and methylation states in parental HCT116 cells. In DKO cells, there were 23,301 hyper-accessible sites, and 3,166 hypo-accessible sites, compared to parental HCT116 cells (Fig. 3A; | log_2_ fold change | > 1, q < 0.01); 16,170 THS sites observed in HCT116 cells were unchanged in DKO cells (| log_2_ fold change | < 1, q > 0.8). Compared to unchanged sites, hyper-accessible sites in DKO cells were depleted of DNA methylation (Fig. 3B); these sites were enriched for ATF3, FOSL1, FOSL2, BATF, AP1 and JUNB binding motifs (Fig. 3C). These TFs were previously shown to interact more strongly to their binding motifs when unmethylated (methyl-minus TFs (Yin et al. 2017)). We infer that chromatin hyper-accessibility at these sites in DKO cells was due to enhanced binding of the methyl-minus TFs when methylation was diminished; this had the effect of limiting nucleosome deposition, thus enabling increased chromatin accessibility. Conversely, hypo-accessible sites in DKO cells were modestly depleted of DNA methylation (Fig. 3B), and enriched for SP1, NFYA, SP5, KLF9, KLF14, and KLF3 binding motifs (Fig. 3D). These TFs were previously shown to exhibit less binding when their sites were unmethylated (methyl-plus TFs (Yin et al. 2017)). We infer that chromatin hypo-accessibility at these sites in DKO cells was due to reduced binding of the methyl-plus TFs when methylation was diminished, and that this led to increased nucleosome deposition, and reduced chromatin accessibility. In support of this is the observation that promoters showing the greatest increases in chromatin accessibility in DKO cells were also the promoter that were most extensively hypomethylated (Fig. 3E). These findings and conclusions are consistent with previously described mechanisms whereby TF binding can regulate nucleosome density (Zaret and Carroll 2011). These conclusions may be tempered by the fact that we are assaying methylation at accessible sequences, the same loci, when in an inaccessible state, are underrepresented in our methylation analyses.

**Figure 3:**
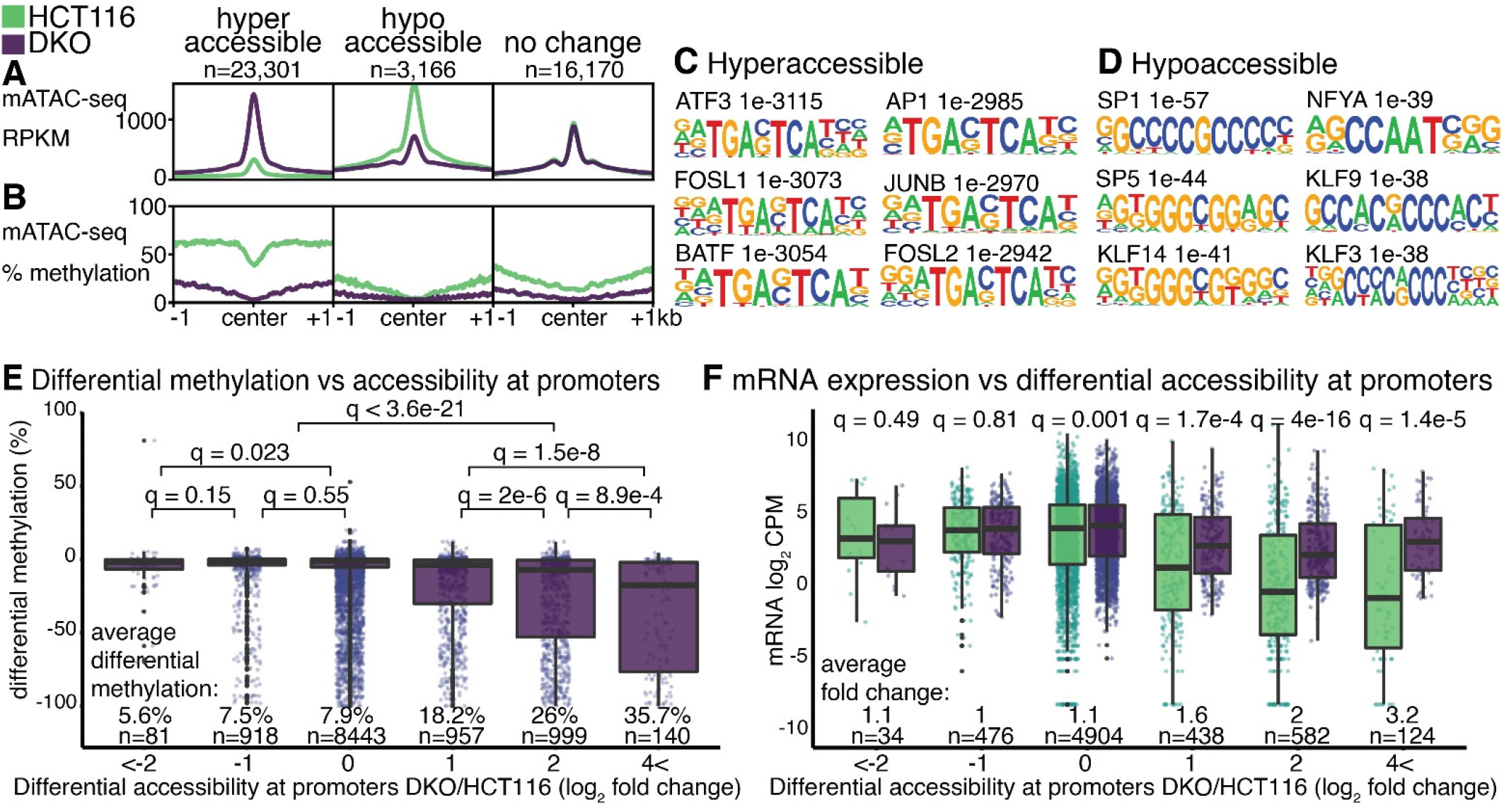
Accessibility and methylation at peaks; (A, B) significantly changed mATAC-seq hyper-accessible (log_2_ fold change > 1, q < 0.01, n = 23,310 peaks), hypo-accessible (log_2_ fold change < −1, q < 0.01, n = 3,166 peaks), and unchanged peaks (| log_2_ fold change | < 1, q > 0.8 n = 16,170). Motifs enriched in (C) hyper- and (D) hypo-accessible sites compared to unchanged sites. (E) DNA methylation changes at promoters binned by accessibility, reported as the change in methylation ratio of DKO cells relative to HCT116 (DKO/HCT116). (F) mRNA expression changes in DKO cells relative to HCT116, reported as log_2_ CPM, at genes binned by differential accessibility of their promoters as in (E). q values are for Wilcoxon tests with Benjamini-Hochberg correction.

To assess how promoter accessibility states detected by mATAC-seq relate to gene expression, we queried existing RNA-seq data from HCT116 and DKO cells (Blattler et al. 2014). Promoters that were hypo-accessible in DKO cells exhibited no significant gene expression changes relative to the corresponding promoters in parental HCT116 cells. At promoters that exhibited no differences in accessibility in the two cell types, there were significant, but very modest differences in mean expression levels. At promoters that were hyper-accessible in DKO cells, we observed substantial and significantly higher levels of expression in DKO cells relative to HCT116, with expression differences increasing as accessibility increased (Fig. 3F). These are in accordance with previous findings (Kelly et al. 2012), further validating the utility of mATAC-seq, and demonstrating the concordance between the extent of chromatin accessibility at promoters, and promoter activity.

Our analyses so far have separately examined methylation and chromatin accessibility results from mATAC-seq. We next combined methylation and accessibility data to take advantage of added value of the combined results afforded by mATAC-seq. We first performed *k*-means clustering of DNA methylation levels at THS sites in HCT116 and DKO cells. DNA methylation at mATAC-seq peaks in HCT116 cells formed five distinct clusters (Fig. 4A). In Cluster 1, accessible peaks, and the 1kb intervals flanking the peaks, were hypermethylated in HCT116 relative to DKO cells, with the flanks exhibiting more hypermethylation. Clusters 2 and 3 were hypomethylated at peak centers in both cells; the clusters were respectively hypermethylated in HCT116 cells in one or the other of the two intervals flanking the peaks. Cluster 4 was hypermethylated over the peaks only in HCT116 cells, and hypomethylated in the peak and flanks in DKO cells. Cluster 5 was hypomethylated in the peaks and flanks of both cell types (Fig. 4A, C, D).

**Figure 4:**
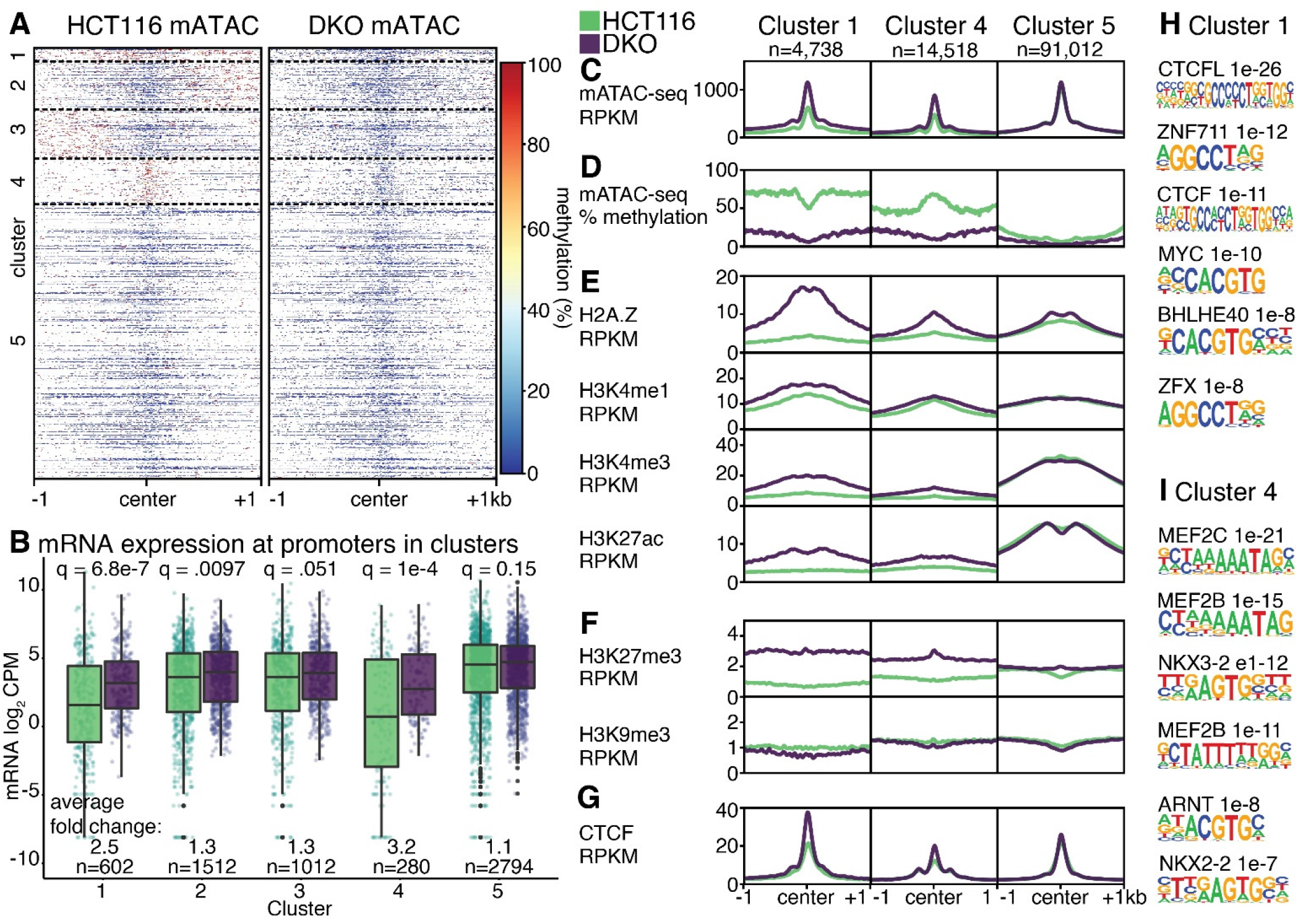
Combined Accessibility and methylation analysis. (A) DNA methylation at mATAC-seq peaks from HCT116 and DKO cells form 5 distinct clusters by DNA methylation. (B) mRNA expression in log_2_ CPM at identified clusters. Features in clusters 1, 4 and 5 are depicted according to (C) accessibility, (D) DNA methylation, (E) activating histone modifications, (F) silencing histone modifications, and (G) CTCF. Motifs enriched in cluster 1 (H), and cluster 4 (I), compared to cluster 5. q values are for Wilcoxon tests with Benjamini-Hochberg correction.

When we assessed mRNA expression from promoters within the five clusters, differences between DKO and HCT116 emerged that varied according to cluster. Promoters in DKO cells from Clusters 1 and 4 were significantly more active than the corresponding promoters from the same clusters in HCT116 cells, with respective increases in mRNA of 2.5-, and 3.2-fold (Fig. 4B). Clusters 2 and 3 exhibited a modest change of 1.3-fold between the cell types. Cluster 5, which was both hypomethylated and hyper-accessible in both cell types, showed no difference in expression.

Besides the differences in DNA methylation and expression, the clusters have additional distinguishing features. There are more promoters, CpG islands, and exons in Cluster 1 compared to Cluster 4; and more intronic and distal intergenic elements in Cluster 4 compared to Cluster 1 (Fig. S3A, B). One feature is the broad domain of H2A.Z in Cluster 1 that accompanied the loss of DNA methylation in DKO cells (Fig. 4E). This finding is consistent with reports that DNA modification and H2A.Z are mutually antagonistic (Zilberman et al. 2008). In Cluster 4, where hypermethylation in HCT116 cells is largely confined to the mATAC-seq peak, there was also an increase in H2A.Z in DKO cells, with the increase being more modest and confined to a narrower portion of the 2kb window displayed. Additional histone modifications associated with active chromatin (H3K4me1, H3K4me3, H3K27ac) were elevated in DKO cells near Cluster 1 mATAC-seq peaks, but these effects were limited or absent in Cluster 4 (Fig. 4E). Like H2A.Z, H3K27me3 was increased in DKO cells at Cluster 1, with the effects also being more modest at Cluster 4 (Fig. 4F). This is also consistent with antagonism reported between H3K27me3 and DNA methylation (Lindroth et al. 2008). In contrast to these histone modifications and variants, H3K9me3 at mATAC peaks was largely unaffected by DNMT-loss. Cluster 5 shows no DNA methylation changes between the two conditions, and there were little to no changes in deposition of histone modifications and variants.

Motifs for TFs, and CTCF binding also varied by cluster. Cluster 1 is enriched for motifs recognized by DNA methyl-plus TFs such as CTCFL, MYC, and BHLHE40; ZFX and ZNF711 contain similar motifs to ZNF704, a methyl-minus transcription factor (Fig 4H). Of the top five TFs enriched in Cluster 4, three are MEF-family TFs, followed by ARNT, which was previously suggested to be methyl-sensitive (Lay et al. 2015) (Fig. 4I). ARNT motifs share substantial sequence identity with BHLHE40, a methyl-minus TF.

## DISCUSSION

ATAC-seq identifies nucleosome-depleted regions of the genome, which are arguably the most relevant for gene regulation within cells. By including bisulfite treatment in the workflow, mATAC-seq targets DNA methylation profiling to open chromatin sites that are enriched for active regulatory regions of the genome. Accordingly, mATAC-seq queries the functional methylome of cells, using relatively few reads compared to WGBS. This is in contrast to other assays for DNA methylation that query the entire genome, or other domains that may not be regulatory.

By applying mATAC-seq to the well-characterized HCT116 cell line, and its DNA methylation-deficient DKO derivative, we demonstrated that mATAC-seq detects DNA methylation patterns that agree with both previously described WGBS results, and with our Omni-ATAC-seq results. These tests validated the fidelity, and compatibility of combining tagmentation and bisulfite treatment steps in the mATAC-seq workflow. DKO cells had many hyper-accessible sites relative to parental HCT116 cells, and these sites exhibited loss of methylation. These same regions were also enriched for methyl-minus TF binding sites, which interact more strongly with DNA when the sites are in an unmethylated context. This highlights the instructive role of TF binding for nucleosome occupancy in the genome. Specifically, our data indicate that when DNA is unmethylated, it facilitates the recruitment of methyl-minus TFs, and that these in turn enable chromatin to assume an open state. Our data also revealed that hyper-accessible and hypomethylated domains in DKO cells were enriched for the histone variant H2A.Z, implicating this factor in limiting DNA methylation, and nucleosome density at sequences where it is recruited. In contrast, regions that displayed no change in methylation showed little change in accessibility. We did not observe a depletion of H3K9me3 at sites with increased accessibility, and decreased DNA methylation, confirming statements in previous studies (Blattler et al. 2014). Such findings were made possible by the combination of DNA methylation, and open chromatin status provided by mATAC-seq.

We envisage that mATAC-seq could be applied to many other systems. For example, HCT116 derivatives carrying single *DNMT* knockouts (Rhee et al. 2000; Rhee et al. 2002) would enable us to identify regulatory elements the different DNMTs individually target for methylation, and their respective influences on nucleosome placement. The various DNMTs have been shown to regulate DNA methylation states by independent, as well as cooperative mechanisms (Liang et al. 2002). Repetitive elements are common targets of the DNMTs, and the resulting DNA methylation contributes to their silencing. However, silencing can occur when DNMT activities are impaired, indicating that compensating mechanisms can silence transposons, likely involving H3K9 methylation, and possibly other chromatin modifications (Horard et al. 2009; Karimi et al. 2011; Walter et al. 2016; Jorda et al. 2017). Querying the specificities of the DNMTs, and their influences on chromatin accessibility at repetitive elements using mATAC-seq can elaborate mechanisms underlying repeat regulation.

Additionally, by using HCT116 DKO cells, we studied the effects of DNA methylation depletion that arose by passive mechanisms due to a lack of DNA methylation maintenance. Active demethylation by TET dioxygenases, and AID/APOBEC deaminases, occurring during differentiation and response to stimuli, is a distinct process. Applying mATAC-seq to stem cell differentiation – including under conditions where these active demethylation mechanisms are altered – can reveal both the combinatorial changes in accessibility and DNA methylation, and the effects active DNA methylation mechanisms have on chromatin state during differentiation.

Pioneer transcription factors have the unique property of binding chromatin that is generally inaccessible to other transcription factors (Zaret and Mango 2016). Application of mATAC-seq to systems where pioneer factor functions are altered can reveal the influences these factors have on both chromatin accessibility, and methylation state of the underlying DNA.

Our protocol for mATAC-seq can potentially be integrated with existing methods for combinatorial detection of other DNA modifications including 5-hydroxymethylcytosine (Yu et al. 2012; Booth et al. 2013), 5-formylcytosine (Song et al. 2013), and 5-carboxycytosine (Wu et al. 2016). ChIPmentation uses Tn5 tagmentation in a chromatin immunoprecipitation workflow (Schmidl et al. 2015). This too could be implemented, using steps we developed for mATAC-seq, to identify locations of DNA-bound proteins, and the underlying DNA modification states in a combinatorial detection strategy similar to other methyl-ChIP strategies (Brinkman et al. 2012; Statham et al. 2012). Combinatorial indexing as a low-cost strategy to query single cells can be used to enable the extension of mATAC-seq to a single cell format; specifically, methods such as single-cell combinatorial indexing assay for transposase accessible chromatin using sequencing (sci-ATAC-seq) (Cusanovich et al. 2015), and for methylation analysis (sci-MET) (Mulqueen et al. 2018). Some alterations are necessary to adapt our mATAC-seq protocol for single cell sequencing, including, extending the indexed adapter set to use methylated sci-ATAC-seq adapters during tagmentation, followed by split-pooling, methylated end-repair, bisulfite conversion in a 96-well format, and PCR. The challenge to this approach is the depletion of reads due to the destructiveness of bisulfite conversion, and the limited sequence complexity in bisulfite converted reads.

## MATERIALS AND METHODS

### Cell Culture

Cultured cells (#28 HCT116 Parental and #343 DKO) were procured from the Genetic Resources Core Facility at Johns Hopkins School of Medicine and cultured in McCoy’s Modified 5A Medium containing 10% heat-inactivated FBS and 1× Penn/Strep (Gibco #15140122). Cells for each experiment were grown apart for at least 2 passages before library preparation.

### Genotyping

DNA from each cell line was extracted using EZ-10 Spin Columns (Bio Basic #BS427) following the manufacturer’s protocol. Genotyping PCR was performed on 50ng genomic DNA using oligos from Table S2 from (Das and Chadwick 2016) for 40 cycles using GoTaq (Promega #M3001) (94°C 2 minutes, 40 cycles of: [94°C 30 seconds, 60°C 30 seconds, 72°C 30 seconds], 72°C 5 minutes) and run on a 2% agarose gel. Cells were confirmed to be Mycoplasma-free and HeLa-free via PCR (Rahbari et al. 2009; Young et al. 2010) on 50ng genomic DNA and cell-culture media (Figure S4).

### Omni-ATAC-seq

Cells were trypsinized and subsequently inactivated in cell culture media. Following inactivation, cells were pelleted and resuspended in cold PBS (without Ca++ and Mg++). Cells were stained with Trypan Blue and counted on a hemocytometer. Lysis and tagmentation were performed exactly as described (Corces et al. 2017) with modifications to inactivation and size selection. Briefly, 100,000 HCT116 Parental and DKO cells were lysed on ice for 3 minutes in 50 μL ice-cold Lysis Buffer (10mM Tris pH 7.4, 10mM NaCl, 3mM MgCl_2_, 0.1% NP-40, 0.1% TWEEN 20, 0.01% Digitonin in DEPC H2O), resuspended in 1mL ice-cold RBS-Wash (10mM Tris pH 7.4, 10mM NaCl, 3mM MgCl_2_, 0.1% TWEEN 20) and pelleted at 4°C at 500 × g for 10 minutes. Tagmentation was performed in 1 × Tagmentation Buffer (10mM Tris pH 7.4, 5mM MgCl_2_, 10% DMF, 33% PBS, 0.1% TWEEN 20, 0.01% Digitonin) using 100nM Tn5 Transposase for 30 minutes at 37°C. Tagmentation was inactivated with the addition of 5 volumes SDS Lysis Buffer (100mM Tris pH 7.4, 50mM NaCl, 10mM EDTA, 0.5% SDS in H_2_O) and 100μg Proteinase K (Invitrogen #25530049) for 30 minutes at 55°C followed by Isopropanol precipitation using GlycoBlue (Invitrogen #AM9516) as a carrier. DNA was size selected using Ampure XP beads (Beckman Coulter # A63880) using a 0.5 × volumes to remove large fragments followed by a 1.8 × final volume according to the manufacturer’s instructions. PCR was performed for using Q5 DNA polymerase (NEB #M0491S) with 1 × GC buffer (72°C 5 minutes, 98C 30 seconds, 11 cycles of: [98°C 10 seconds, 65°C 30 seconds, 72°C 30 seconds], 72°C 5 minutes) followed by a final cleanup using a 1.8 × volumes of Ampure XP beads according to the manufacturer’s instructions.

### methylATAC-seq

Cell lysis was performed identically to Omni-ATAC-seq. Tagmentation was performed on 250,000 HCT116 Parental and DKO cells using 700nM Tn5 Transposase assembled using preannealed Tn5ME-A_mC and Tn5ME-B_mC (Table S3) for 30 minutes at 37°C following the addition of 0.01ng of unmethylated Lambda DNA (Promega #D1521). We recommend performing a titration of Tn5 transposase to nuclei input to assay minimum amounts required as in Figure S5. Inactivation and size-selection were performed identically to our modified Omni-ATAC-seq protocol. Tagmented DNA was End-Repaired for 30 minutes at 37°C (5U Klenow Exo-(NEB #M2012S), 1 × NEB Buffer 2, and 0.5 mM/each dATP, dGTP, dTTP, and 5-mdCTP (NEB #N0365S)) similar to T-WGBS (Lu et al. 2015) and X-WGBS (Suzuki et al. 2018). End repair was cleaned using a 1.8 × volumes of Ampure XP beads according to the manufacturer’s instructions. 10% of the product was kept for quality control PCR (Fig. S6A). Bisulfite conversion was performed using EZ DNA Methylation-Lightning (Zymo #D5030T) following the manufacturer’s protocol. PCR was immediately performed using PfuTurbo Cx (Agilent #600410) (94°C 2 minutes, 13 cycles of: [98°C 10 seconds, 6°5C 30 seconds, 72°C 30 seconds], 72°C 5 minutes) (Fig S6B) followed by a final cleanup using a 1.8 × volumes of Ampure XP beads according to the manufacturer’s instructions.

### Tn5 Transposase

Tn5 was produced exactly as described (Picelli et al. 2014) with no modifications. For Omni-ATAC-seq, Tn5 transposase was assembled as described (Adey and Shendure 2012) using pre-Annealed Tn5MEDS-A and Tn5MEDS-B from Table S3. For methylATAC-seq, Tn5 transposase was assembled using pre-Annealed Tn5ME-A_5mC and Tn5MEB_5mC oligonucleotides from Table S3. Oligonucleotides were annealed by combining ME-A or ME-B oligos to Tn5MErev and incubating for 2 minutes at 94°C followed by a 0.1°C/s ramp to 25°C. Enzyme was stored at −80°C.

### Data Analysis

Libraries were quantified using the Qubit dsDNA HS Assay Kit (ThermoFisher #Q32854). High-throughput sequencing was performed by the Cornell University Genomics Facility on the Illumina NextSeq 500 with single-end 75bp reads. Trimming for mATAC-seq and Omni-ATAC-seq was performed using fastp (Chen et al. 2018) -q 20 -l 20 -a CTGTCTCTTATACACATCT. Trimming for ChIP-seq, RNA-seq, and WGBS data was performed using fastp -q 20 -l 20 -a AGATCGGAAGAGCACACGTCTGAACTCCAGTCAC.

#### Alignment to hg19

In this study we used GRCh37 instead of GRCh38 to match previous studies using similar cells and methods. These results would not be affected by this change because we do not study centromeric sequences and predominantly discuss changes at promoters.

#### ChIP-seq hg19

Trimmed FASTQ files were aligned using BWA-MEM (Li and Durbin 2010) to hg19. Reads were deduplicated using Picard [http://broadinstitute.github.io/picard/] MarkDuplicates. HCT116 and DKO ChIP-seq data for H2A.Z, H3K4me3, H3K4me1, H3K27Ac, H3K27me3, H3K9me3, and H3K36me3 data (Lay et al. 2015) were downloaded from NCBI GEO database accession GSE58638. HCT116 and DKO ChIP-seq data for CTCF (Maurano et al. 2015) were downloaded from NCBI GEO database accession GSE50610.

#### RNA-seq hg19

Pair-end trimmed FASTQ files were aligned using HISAT2 (Kim et al. 2015) to hg19. HCT116 and DKO RNA-seq data (Blattler et al. 2014) were downloaded from NCBI GEO database accessions GSE52429 and GSE60106, respectively.

#### Omni-ATAC and mATAC-Seq

Trimmed FASTQ files were aligned using Bismark (Krueger and Andrews 2011) v0.19.0 to hg19 using the following settings: --score_min L,0,-0.6. Bisulfite reads to be used for MethylKit were filtered for non-conversion using Bismark’s filter_non_conversion and deduplicated using deduplicate_bismark. Methylation was extracted using Bismark’s methylation extractor --gzip --bedgraph --counts --ignore 9 --ignore_3prime 9. Reads used for peak calling and ATAC-seq visualization were deduplicated using deduplicate_bismark without filtering for non-conversion. Conversion rate (Table S1) was measured by aligning to the lambda genome (GenBank: J02459.1) and filtered as above; percent conversion rate was calculated as (1-(Total methylated C’s in all contexts)/(Total number of C’s analyzed)) × 100.

#### WGBS hg19

Trimmed FASTQ files were aligned using Bismark v0.19.0 to hg19 using the following settings: --score_min L,0,-0.6. Bisulfite reads to be used for MethylKit were filtered for non-conversion using Bismark’s filter_non_conversion and deduplicated using deduplicate_bismark. Methylation was extracted using Bismark’s methylation extractor --gzip -- bedgraph --counts. HCT116 and DKO WGBS data (Blattler et al. 2014) were downloaded from NCBI GEO database accession GSE60106.

#### Methylation

Differential methylation was quantified using MethylKit (Akalin et al. 2012) at merged HCT116 and DKO mATAC-seq peaks extended to 1kb tiles covering at least 3 CpGs. Promoters were defined as being within 1kb of a TSS using Genomation (Akalin et al. 2015).

#### Peak calling

ATAC-seq peaks were called using HOMER (Heinz et al. 2010) findPeaks localSize 50000 -size 150 -minDist 50 –fragLength 0 -style dnase. ChIP-seq peaks were called using HOMER findPeaks -style histone. CTCF ChIP-seq peaks were called using HOMER findPeaks -style factor. Reads were assigned to peaks merged from HCT116 and DKO cells using featurecounts (Liao et al. 2013) on reads filtered for a minimum log_2_ CPM of 0.5 in at least 2 samples. Differential accessibility was called using DESeq2 (Love et al. 2014) lfcShrink. Hyper- and hypo-accessible peaks were defined as having a | log_2_ FC | > 1 with an adjusted p value < 0.01 in DKO compared to HCT116 parental cells. Promoters were defined as being within 1kb of a TSS using Genomation. FRiP scores in Table S1 and sample correlation in Fig. S2 were quantified using DiffBind (Stark and Brown 2018) on libraries downsampled to 5M reads using Picard DownsampleSam using peaks called by HOMER. Peak overlaps for Fig. 1A and Fig. S2E were generated using ChIPpeakAnno (Zhu et al. 2010). Feature overlaps for Fig. 1B and S3A were generated using ChIPseeker (Yu et al. 2015). Motif enriched in changed peaks were called using HOMER findMotifsGenome to the hg19 genome using unchanged peaks as background.

#### RNA-seq quantification

Unstranded hg19-aligned reads were assigned to hg19 genes using featurecounts inbuilt reference using default settings. Differential expression was quantified using DESeq2 lfcShrink on reads filtered for a minimum CPM of 0.5 in at least 2 samples.

#### Genome browser visualizations

ATAC-seq and mATAC-seq bigWig files were made using eepTools (Ramirez et al. 2016) bamCoverage --binSize 1 --normalizeUsing RPKM -- ignoreForNormalization chrM --scaleFactor N and viewed on UCSC genome browser. ChIP-Seq bigWigs were made using deepTools bamCoverage --binSize 10 --normalizeUsing RPKM -- ignoreForNormalization chrM --scaleFactor N. Scale factor was determined by coverage of peaks called by HOMER shared between HCT116 and DKO via bedops --intersect where N = (% reads in shared peaks in HCT116)/(% reads in shared peaks in DKO) when N > 1.1. Scaling was applied to the following samples: DKO_mATAC 1.877, DKO_H3K27ac_R2 1.48, H3K4me3_R2 = 1.47.

Gene body heatmaps were produced using deepTools plotheatmap --beforeRegionStartLength 2000 --regionBodyLength 3000 --afterRegionStartLength 2000 to Ensembl hg19 APPRIS PRINCIPALS flagged transcripts (Rodriguez et al. 2013). Heatmaps for differential peaks were centered on peaks called by HOMER. Peaks from HCT116 and DKO were combined using BEDOPS for clustering of DNA methylation at THS sites. Clustering was performed using deepTools plotheatmap --kmeans 5, the output and order of which was used for all subsequent heatmaps.

### Data Access

All raw and processed sequencing data generated in this study have been submitted to the NCBI Gene Expression Omnibus (GEO; http://www.ncbi.nlm.nih.gov/geo/) under accession number GSE126215.

## ACKNOWLEDGEMENTS

Jennifer D. Mosher, Ann E. Tate, Jeff C. Mattison, and Peter A. Schweitzer from the Cornell University Biotechnology Resource Center (BRC) for genomic sequencing; Bert Vogelstein for HCT116 cells and derivatives; Önder Kartal, and Erin Chu for proofreading and helpful comments; the Cornell College of Veterinary Medicine and the National Institutes of Health for funding (R01HG006850 and R01GM105243).

