## Supplemental Figure and Tables for "methyl-ATAC-seq measures DNA methylation at accessible chromatin"

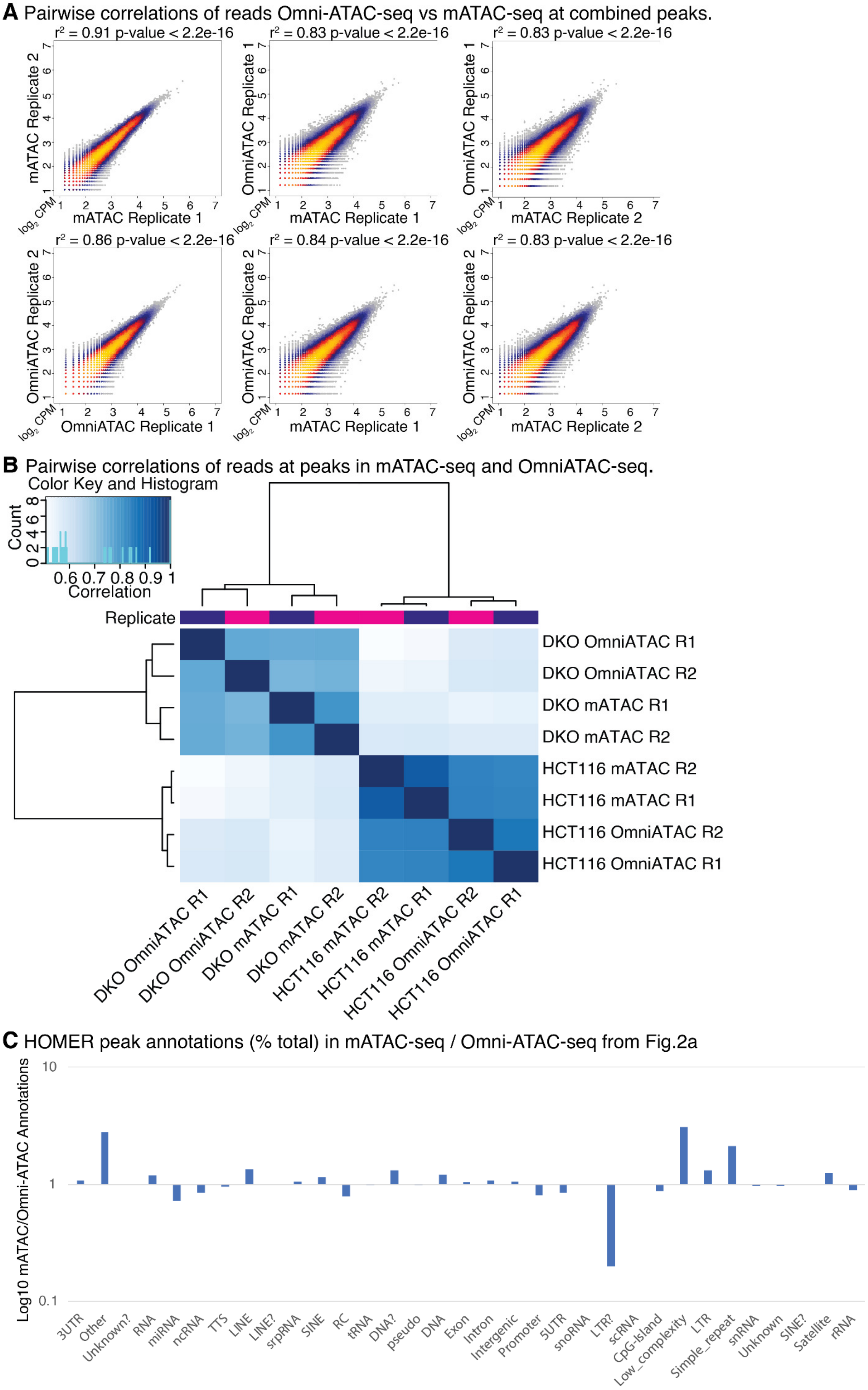
A Pairwise correlations of reads Omni-ATAC-seq vs mATAC-seq at combined peaks.
r² = 0.91 p-value < 2.2e-16
r² = 0.83 p-value < 2.2e-16
r² = 0.83 p-value < 2.2e-16
mATAC Replicate 2
OmniATAC Replicate 1
OmniATAC Replicate 1
mATAC Replicate 1
mATAC Replicate 1
mATAC Replicate 2
log2 CPM
r² = 0.86 p-value < 2.2e-16
r² = 0.84 p-value < 2.2e-16
r² = 0.83 p-value < 2.2e-16
OmniATAC Replicate 2
OmniATAC Replicate 2
OmniATAC Replicate 2
OmniATAC Replicate 1
mATAC Replicate 1
mATAC Replicate 2
B Pairwise correlations of reads at peaks in mATAC-seq and OmniATAC-seq.
Color Key and Histogram
Count
Correlation
0.6 0.7 0.8 0.9 1
Replicate
DKO OmniATAC R1
DKO OmniATAC R2
DKO mATAC R1
DKO mATAC R2
HCT116 mATAC R2
HCT116 mATAC R1
HCT116 OmniATAC R2
HCT116 OmniATAC R1
C HOMER peak annotations (% total) in mATAC-seq / Omni-ATAC-seq from Fig.2a
Log10 mATAC/Omni-ATAC Annotations
10
1
0.1
3UTR
Other
Unknown?
RNA
miRNA
ncRNA
TTS
LINE
LINE?
srpRNA
SINE
RC
tRNA
DNA?
pseudo
DNA
Exon
Intron
Intergenic
Promoter
5UTR
snoRNA
LTR?
scRNA
CpG-Island
Low_complexity
LTR
Simple_repeat
snRNA
Unknown
SINE?
Satellite
rRNA

**Figure S1:** (A) Scatterplots of HCT116 mATAC-seq reads at peaks in libraries downsampled to 5M reads. (B) Pairwise Pearson correlations between mATAC-seq and Omni-ATAC-seq of HCT116 and *DNMT1*/*DNMT3b* Double Knockout (DKO) cells in downsampled libraries. (C) Annotations from peaks (% total) in Omni-ATAC-seq/mATAC-seq in Fig 2A.

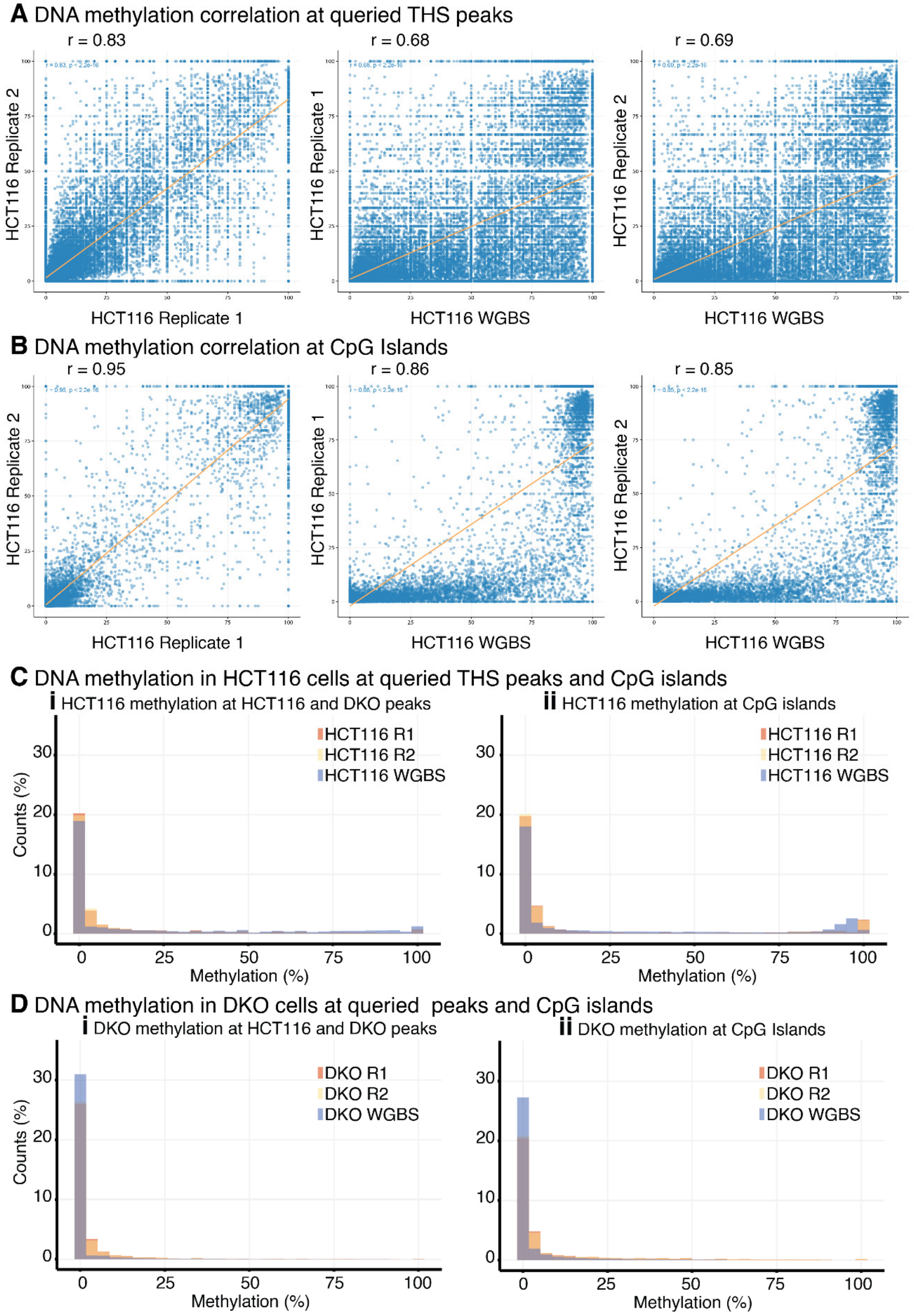


**Figure S2:** Scatterplots of HCT116 mATAC-seq libraries vs WGBS at (A) THS sites and (B) CpG Islands. DNA methylation in mATAC-seq and WGBS in (C) HCT116 at i. merged HCT116 and DKO peaks and ii. CpG islands and (D) DKO at i. merged HCT116 and DKO peaks and ii. CpG islands. Grey bars denote an overlap of all labeled samples, orange bars denote an overlap of mATAC-seq Replicate1 and Replicate2 samples.

**A** Genomic features at THS peaks in clusters

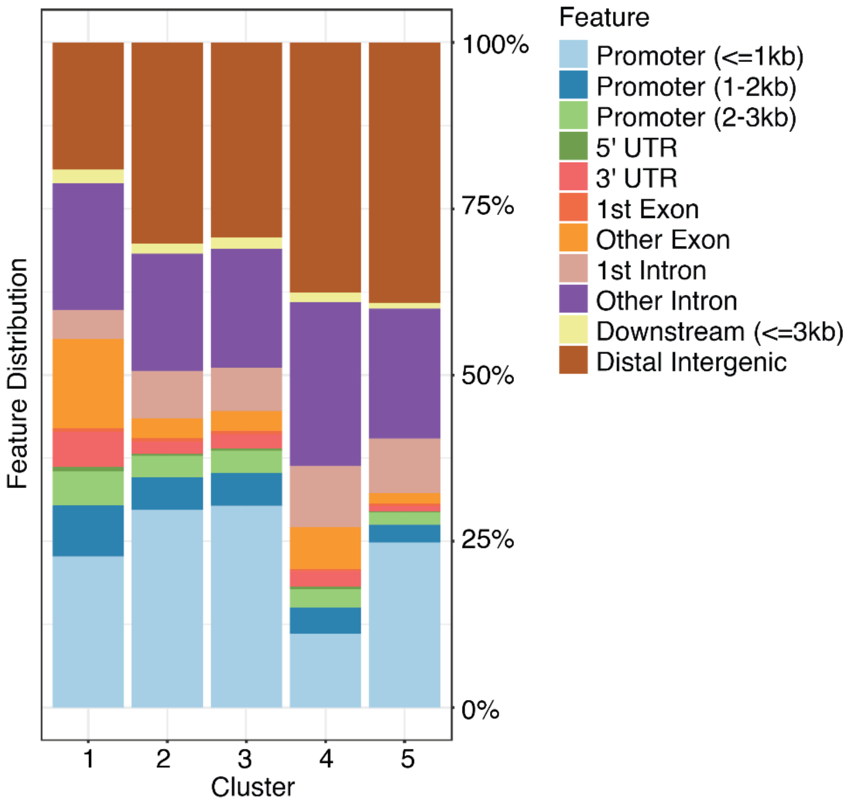


**B** HOMER annotations at Clusters

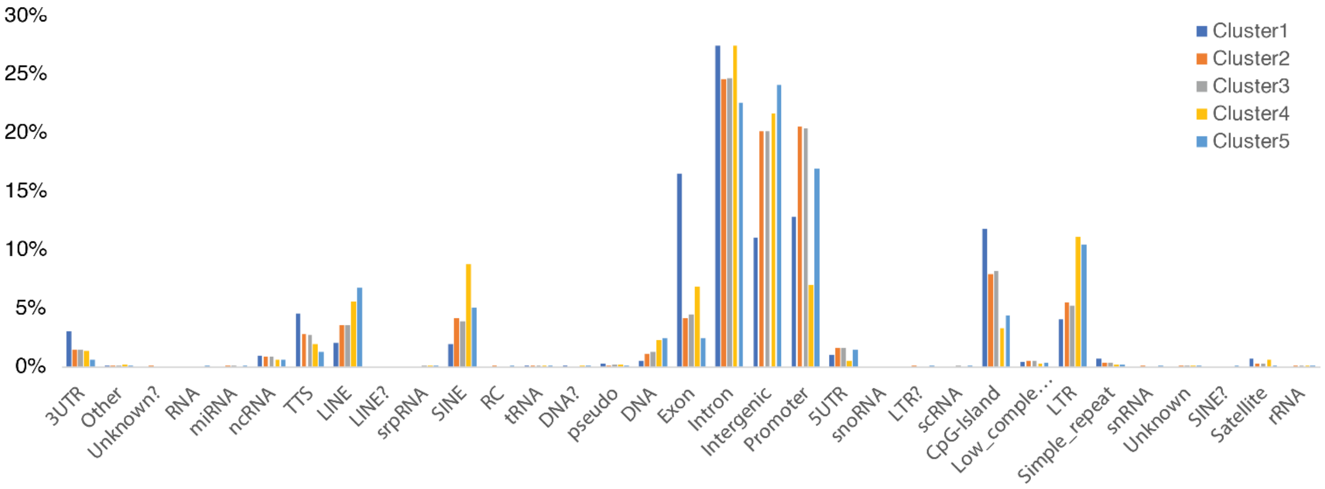


**Figure S3:** (A) Genomic features, and (B) HOMER annotations for clusters shown in Fig. 4.

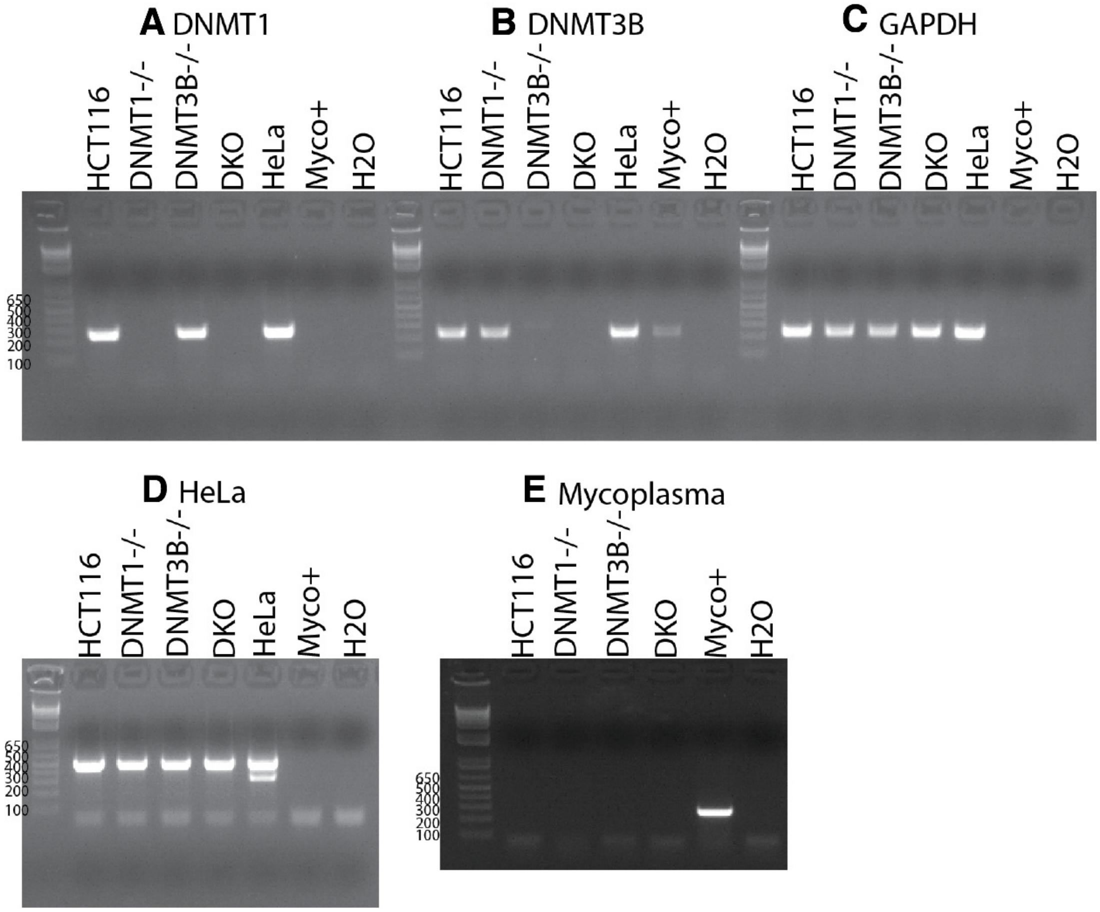


**Figure S4:** Genotyping by PCR to test for presence of *DNMT1* (A), *DNMT3B* (B), *GAPDH* (C), HeLa cell contamination (D), and Mycoplasma (E) in cells used for this study.

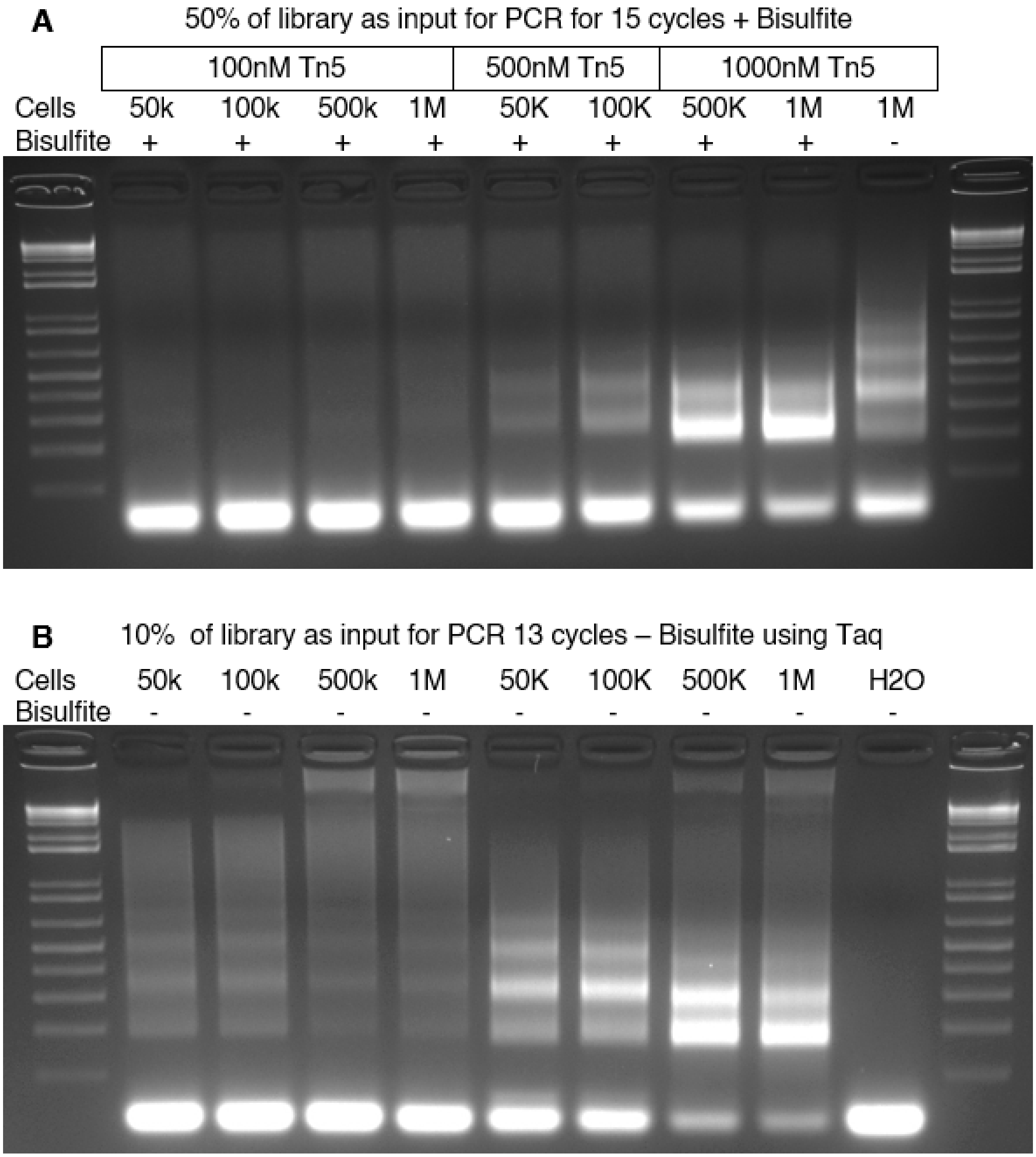


**Figure S5:** Titration of Tn5 transposase compared to input nuclei. An input range of nuclei and Tn5 were tested using libraries either (A) Bisulfite converted library amplified with PfuTurbo Cx (B) not converted library amplified with Taq to assay viable product for sequencing. Samples were run using a 3.5% Agarose gel in TAE; samples were resuspended in loading buffer containing 1X SYBR Gold.

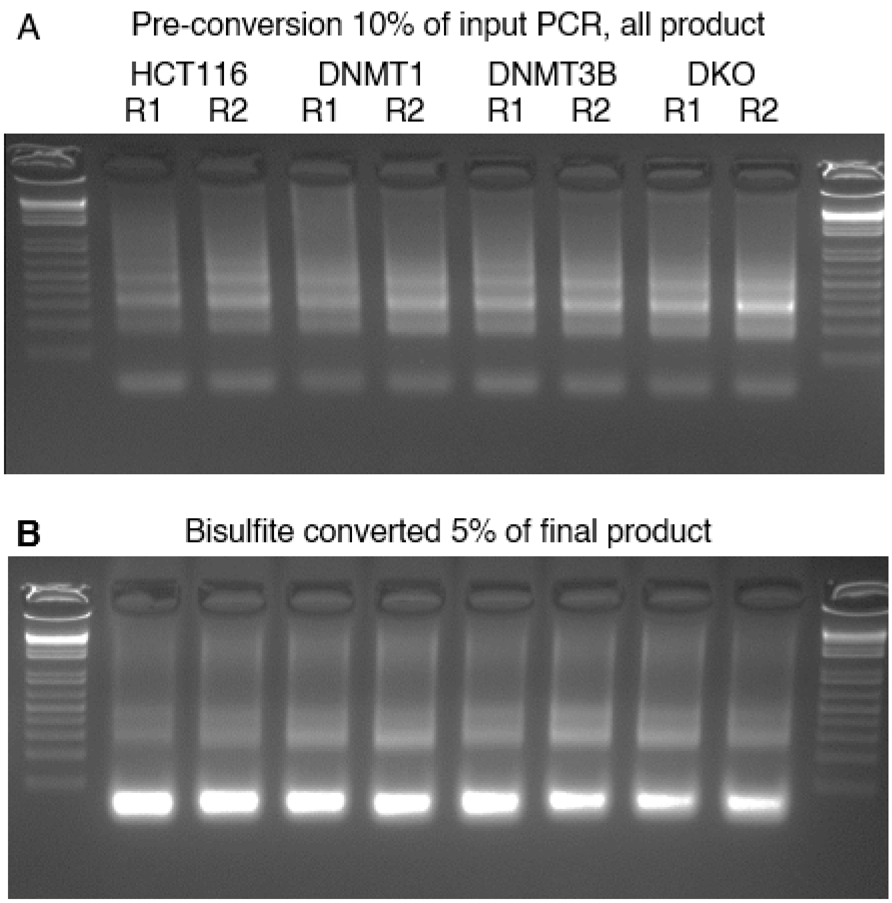


**Figure S6:** QC of final methyl-ATAC-seq libraries. PCR was performed on (A) 10% of pre-converted libraries using Taq (B) bisulfite converted libraries using PfuTurbo Cx. Samples were run using a 3.5% Agarose gel in TAE; samples were resuspended in loading buffer containing 1X SYBR Gold.

| Sample: | Trimmed sequences analyzed (M): | mapping: | duplication: | Conversion rate (Lambda mC/TotalC): | FRiP (5M reads): |
|---|---|---|---|---|---|
| mATAC-seq HCT116 Replicate 1 | 32.6 | 45.40% | 63.73% | 99.76% | 0.45 |
| mATAC-seq HCT116 Replicate 2 | 25.1 | 46.60% | 38.42% | 99.54% | 0.49 |
| mATAC-seq DKO Replicate 1 | 24.6 | 44.70% | 25.12% | 98.93% | 0.31 |
| mATAC-seq DKO Replicate 2 | 23.2 | 46.30% | 26.22% | 99.35% | 0.27 |
| Omni-ATAC-seq HCT116 Replicate 1 | 15 | 89.30% | 17.66% | | 0.26 |
| Omni-ATAC-seq HCT116 Replicate 2 | 14.3 | 89.20% | 18.61% | | 0.28 |
| Omni-ATAC-seq DKO Replicate 1 | 16.5 | 88.50% | 18.92% | | 0.27 |
| Omni-ATAC-seq DKO Replicate 2 | 22.7 | 88.20% | 12.84% | | 0.15 |

**Table S1:** Samples sequenced in this study. Values show millions (M) trimmed sequences analyzed, mapping efficiency using Bismark, conversion rate of unmethylated Lambda-DNA spike-in after filtering, and Fraction of Reads in Peaks (FRiP) score in down-sampled libraries.

| Genotyping oligonucleotides: | |
|---|---|
| DNMT1-Gen-F1: | AAACTGGCAGGTGCTAACTG |
| DNMT1-Gen-R1: | AGATGTGATGGTGGTTTGCC |
| DNMT3B-Gen-F1: | TTGGTTTTGCTCAGAGCCAG |
| DNMT3B-Gen-R1: | ACGTGTGGGCAAGAGATTTC |
| GAPDH-Gen-F1: | GAAGGTGAAGGTCGGAGTC |
| GAPDH-Gen-R1: | GAAGATGGTGATGGGATTTC |
| GPO-3 | GGGAGCAAACAGGATTAGATACCCT |
| MGSO: | TGCACCATCTGTCACTCTGTTAACCTC |
| VM164A: | TGCCTCCTAGATCGTATTCCC |
| VM-164B: | GCACTCTGTGGCATGAAGGT |
| RB164K2: | TGGCTCCTCCCTCTATTATCG |

**Table S2:** Oligonucleotides used in Figure S4 to genotype and verify cell lines were ordered from Integrated DNA Technologies and purified by standard desalting.

| Tn5 oligos: | |
|---|---|
| Tn5MErev: | /5Phos/CTGTCTCTTATACACATCT |
| Tn5ME-A: | TCGTCGGCAGCGTCAGATGTGTATAAGAGACAG |
| Tn5ME-B: | GTCTCGTGGGCTCGGAGATGTGTATAAGAGACAG |
| Tn5ME-A_mC: | T/iMe-dC/GT/iMe-dC/GG/iMe-dC/AG/iMe-dC/GT/iMe-dC/AGATGTGTATAAGAGA/iMe-dC/AG |
| Tn5ME-B_mC: | GT/iMe-dC/T/iMe-dC/GTGGG/iMe-dC/T/iMe-dC/GGAGATGTGTATAAGAGA/iMe-dC/AG |
| i7 Primers: | CAAGCAGAAGACGGCATACGAGAT[i7]GTCTCGTGGGCTCGG |
| i5 Primers: | AATGATACGGCGACCACCGAGATCTACAC[i5]TCGTCGGCAGCGTC |

**Table S3:** Oligonucleotides used to assemble Transposomes and PCR libraries were ordered from Integrated DNA Technologies and purified by standard desalting. '/iMe-dC/' represents an internal methylated dC. '/5Phos/' represents a 5' phosphate. '[i7]' represents Nextera i7 barcodes. '[i5]' represents Nextera i5 barcodes.
